# Identification of a Novel Picorna-like Virus in Coconut Rhinoceros Beetles (*Oryctes rhinoceros*)

**DOI:** 10.1101/2020.04.15.042309

**Authors:** Kayvan Etebari, Matan Shelomi, Michael J. Furlong

## Abstract

A novel Picorna-like virus, tentatively named Oryctes rhinoceros Picorna-like virus 1 (OrPV1), was identified in coconut rhinoceros beetle (*Oryctes rhinoceros*) larvae in Taiwan. The complete genome sequence consisted of 9,665 nucleotides with a polyA tail and included one open reading frame. Conserved structural domains such as Picornavirus capsid protein, RNA helicase, Peptidase and RNA-dependent RNA polymerase (RdRp) were identified through Pfam domain searches. The genome shares approximately 28% identity with other unclassified picornavirals that infect honey bees (Darwin bee virus 2, Bundaberg bee virus 5, and Sacbrood virus) and a recently reported virus from Asian lady beetle (Harmonia axyridis virus 1). We did not detect this virus in any other geographical populations of *O. rhinoceros* collected from the South Pacific Islands and the Philippines. Analysis of the deduced RdRp amino acid sequences showed that the virus clustered with other Picorna-like viruses and separated from other members of family Dicistroviridae and Iflaviridae.

---

The coconut rhinoceros beetle (*Oryctes rhinoceros*) is a pest of palm trees and a serious threat to livelihoods in tropical Asia and the Pacific Islands. New incursions of the pest into previously *O. rhinoceros*-free countries and territories have been reported in recent years, beginning in Guam in 2007 and followed by Hawaii (2013), Solomon Islands (2015) and most recently Vanuatu and New Caledonia in 2019 (Etebari et al., 2020). It has been suggested that this resurgence and spread of the pest is related to an Oryctes rhinoceros nudivirus (OrNV) resistant population, “CRB-G”, which is identified by single nucleotide polymorphisms (*snp*s) in the Cytochrome C Oxidase subunit I (COI) (Marshall et al., 2017; Reil et al., 2018). The OrNV was isolated from Malaysian populations of *O. rhinoceros* in the early 1960s and its successful establishment, first in Samoa and then elsewhere, reduced pest damage to palms in the region (Bedford, 2013; Huger, 2005). Currently, there is no information explaining why CRB-G, this newly named *O. rhinoceros* haplotype, is not susceptible to the OrNV that is so effective against other geographical haplotypes already established in the Pacific. To date, there is no report on viral diversity in different haplotypes of *O. rhinoceros*. Identification of the viral community associated with *O. rhinoceros* will provide new insights into the potential interaction of insect specific viruses and their hosts and will contribute to an improvement of current biological control programs.

In a previous study (Shelomi et al., 2019), *O. rhinoceros* larvae were collected from decaying coconut (*Cocos nucifera* L.) logs in public land in Jiuru Township, Pingtung County, Taiwan. The total RNA was separately extracted from the fat bodies, hindguts, midguts, and gastric caecae of four late-instar larvae. RNA quality was measured with a NanoDrop™ spectrophotometer and sent to the National Taiwan University TechComm Next Generation Sequencing Core for RNA library construction (mRNA polyA base) and sequencing (Illumina HiSeq 4000, paired end 150 bp). Raw data is available at the NCBI Short Reads Archive, Accession Numbers SRX5979158-61 (Shelomi et al., 2019).

The CLC Genomics Workbench version 12.0.1 was used for bioinformatics analyses. All libraries were trimmed from any vector or adapter sequences remaining. Low quality reads (quality score below 0.05) and reads with more than two ambiguous nucleotides were discarded. To confirm the mitochondrial lineage of *O. rhinoceros* individuals, the trimmed reads were mapped to *O. rhinoceros* COXI gene sequences (MN809502-MN809525) and OrNV complete genome sequence (MN623374). All *O. rhinoceros* individuals in these RNAseq libraries were free from OrNV and belonged to the mitochondrial lineage CRB-G that was previously reported as resistant to OrNV.

To identify novel viruses, all reads were mapped back to the *O. rhinoceros* draft genome (unpublished data) and unmapped reads were retained for *de novo* assembly. The contigs were constructed with kmer size 45, bubble size 50 and minimum length of 500 bp. The contigs were corrected by mapping all reads against the assembled sequences (min. length fraction=0.9, maximum mismatches=2). The generated contigs were compared to the NCBI viral database using local BLAST and BLASTx algorithms. We set the e-value to 1×10-10 to maintain high sensitivity and a low false-positive rate. To detect highly divergent viruses, we also performed domain-based searches by comparing the assembled contigs against the Conserved Domain Database (CDD) version 3.14 and Pfam v32 with an expected value threshold of 1×10−3. Sequences with positive hits to virus polymerase (RNA-dependent RNA polymerase (RdRp) domain: cd01699) were retained.

The Oryctes rhinoceros Picorna-like virus 1 (OrPV1) sequence comprises a single positive-sense single-stranded RNA genome of 9,665 nucleotides, with a natural 3’ poly-A tail and 3’ untranslated region (3’UTR), a short 5’ UTR, and one open reading frame (ORF) of 8,844 nucleotides (Figure 1). A single polyprotein structure with several conserved domains such as Picornavirus capsid protein, RNA helicase, Peptidase and RNA-dependent RNA polymerase (RdRp) was predicted, which is consistent with the genome arrangement of other viruses within the order Picornavirales (Table 1). The viral genomic sequence was found to be A+U rich (64.1%) and it has a predicted molecular mass of 363.477 kDa and theoretical isoelectric point (pI) of 6.58. We found three motifs of Hel-A (GX2GXGKS), Hel-B (QX5DD) and Hel-C (KX5PX5CSN), which are present in the helicase domain of picorna-like viruses, in the helicase region of OrPV1 (Figure 2). However, there is little difference in Hel-C among picornavirales, which previously spoted in small brown planthopper iflavirus (Wu et al., 2019). No large ORF was found in the inverse orientation of the OrPV1 genome, suggesting that this is a positive-strand RNA virus. The OrPV1 genome was built by 28,853 assembled reads that showed higher coverage towards the RdRp region and 3’ ends. The annotated genomic sequence of this virus has been deposited in GenBank under the accession number MT334824.

**Table 1.** The identified conserved domains on OrPV1 polyprotein sequence.

| Domain ID | Code | Description | Start-End | E-value |
| --- | --- | --- | --- | --- |
| RdRp | cd01699 | RNA-dependent RNA polymerase | 2588-2886 | 1.50E-54 |
| RdRp | pfam00680 | RNA-dependent RNA polymerase | 2451-2919 | 4.13E-54 |
| RNA_helicase | pfam00910 | RNA helicase | 1346-1452 | 6.68E-19 |
| rhv_like | cd00205 | Picornavirus capsid protein domain-like | 530-678 | 7.45E-11 |
| rhv_like | cd00205 | Picornavirus capsid protein domain-like | 238-374 | 1.73E-10 |
| rhv_like | cd00205 | Picornavirus capsid protein domain-like | 777-956 | 1.85E-07 |
| Peptidase_C3G | pfam12381 | Tungro spherical virus-type peptidase (Protease) | 2231-2404 | 8.75E-06 |

**Figure 1.**
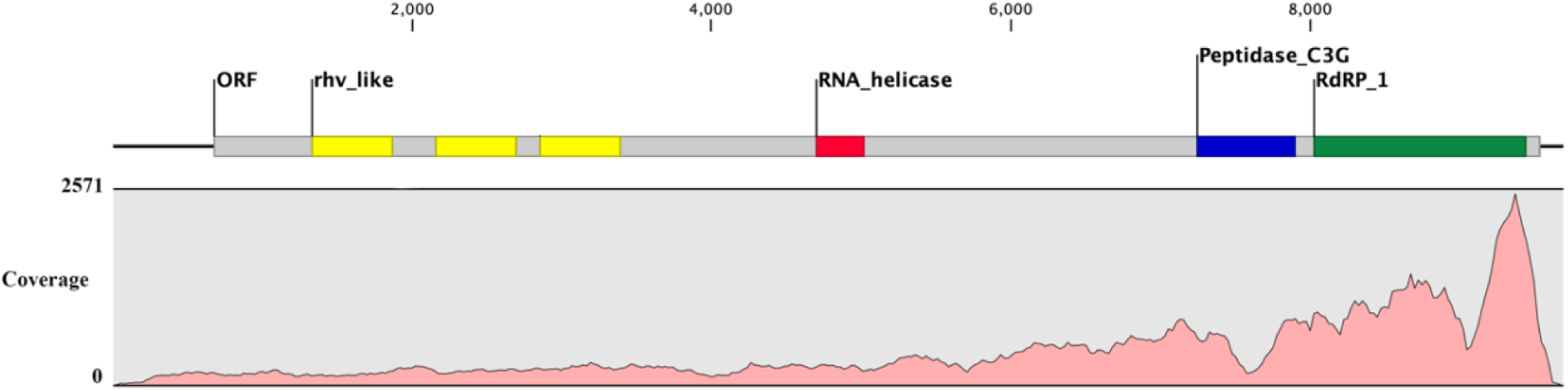
Schematic presentation of the genome of OrPV1. The location of predicted conserved domains is in the upper panel. The reads were mapped back to the sequenced genome to display the coverage and sequence depth (lower panel).

**Figure 2.**
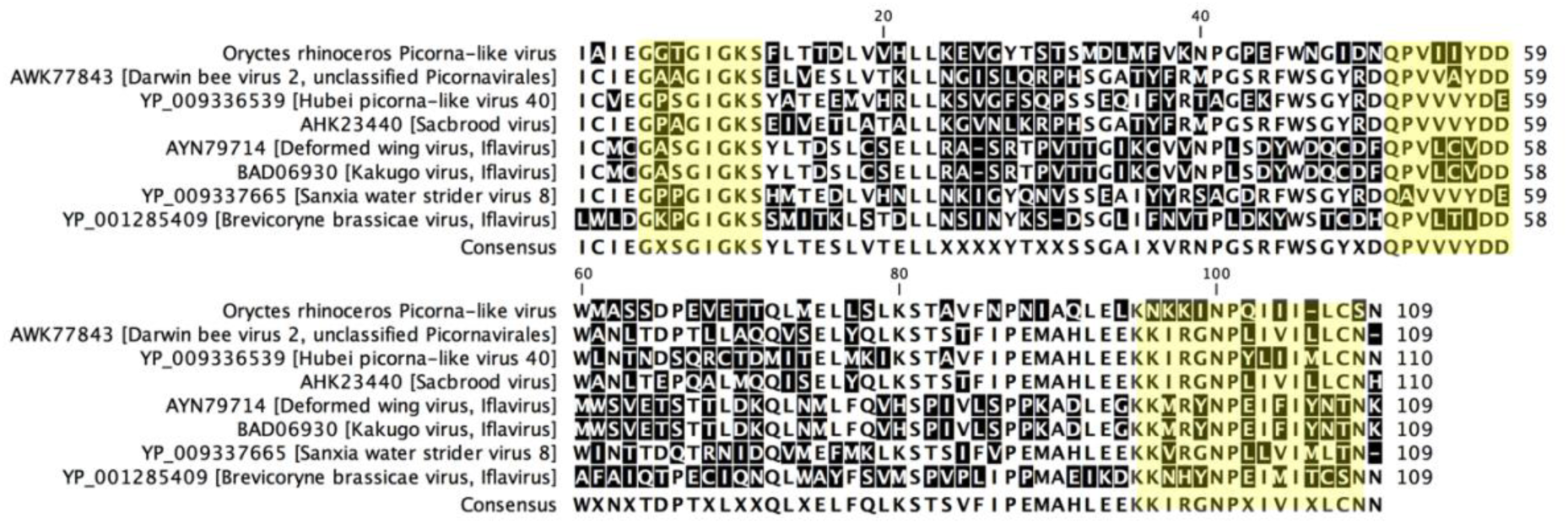
Deduced amino acid sequence alignment for helicases of OrPV1 and some selected Picornavirales. The three conserved motifs Hel-A, -B and -C are highlighted in yellow. The numbers above the sequences indicate the location on helicase domain.

This virus was identified in three libraries (gastric caeca, mid-gut and hind-gut tissues) but no virus sequence was detected in fat bodies. A recently discovered picorna-like virus in *Helicoverpa armigera* (HaNv), is typically found only infecting gut tissue of both adults and larvae (Yang et al., 2019). OrPV1 may similarly be specific to gut tissue. To determine if OrPV1 was associated with a particular geographical region, we looked for this virus in other *O. rhinoceros* RNAseq libraries that we recently generated from beetle gut tissues from the Solomon Islands, Papua New Guinea, Fiji and the Philippines, but no virus sequence has been found so far.

The deduced amino acid sequence of predicted RdRp regions of several viruses from the order Picornavirales were obtained from GenBank to create a maximum likelihood phylogeny tree. The protein distance was measured by the Jukes-Cantor algorithm to construct the tree topologies and its statistical significance was calculated based on 500 bootstraps. The maximum likelihood phylogeny separated the viruses into three major clades belonging to the families Dicistroviridae, Iflaviridae and unclassified Picornavirales or Picorna-like viruses (Figure 3). The NCBI blast search indicated that the complete genome of OrPV1 has around 28% shared identity with other unclassified picornavirales that infect honey bees (*Apis mellifera*) including Darwin bee virus 2, Bundaberg bee virus 5, and Sacbrood virus, as well as a recently reported virus from an Asian lady beetle (Harmonia axyridis virus 1) isolated in China.

**Figure 3.**
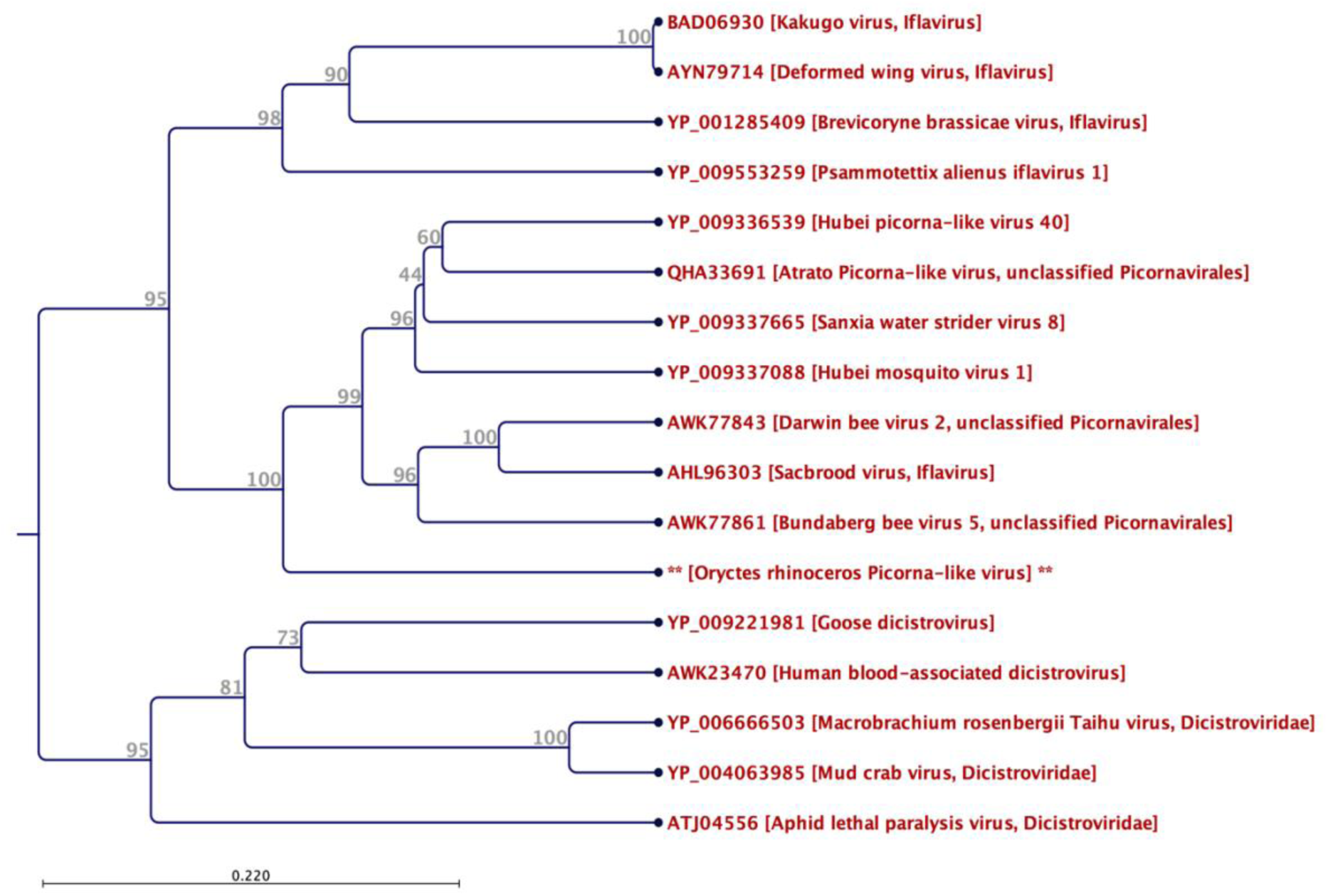
Maximum likelihood phylogeny based on the deduced amino acid sequence of RdRp region of selected Picornavirales. The novel Oryctes rhinoceros picorna-like virus (OrPV1) has been grouped with other unclassified Picornavirales and Separated from members of Dicistroviridae and Iflaviridae families.

Members of three positive-sense single-stranded RNA virus families such as Dicistroviridae (Bonning and Miller, 2010), Iflaviridae (Valles et al., 2017) and Picornaviridae (Zell, 2018) are widely reported from insects. These families were included in a new virus order Picornavirales (Le Gall et al., 2008). Many newly discovered insect viruses with small isometric virions are assigned under the supergroup picorna-like viruses, which are not yet classified by the International Committee on Taxonomy of Viruses (ICTV). Insect-specific small viruses can become an important component of pest management, but the potential that they offer has received little recognition in a pest control landscape dominated by a focus on baculoviruses (arthropod-specific DNA viruses). Recently, some members of small RNA viruses have shown the potential to be used as biopesticides to control insects such as aphids, fire ants, moths and vectors of human diseases (Chen et al., 2012; Valles and Hashimoto, 2009).

Infection by several picorna-like viruses in vertebrate hosts is well characterized, but our knowledge about those infecting invertebrates is very limited (Ryabov, 2017). Most of these insect small RNA viruses do not cause acute infections in their host, but their persistent non-symptomatic infections can interact with large DNA viruses that are employed as biological control agents. For example, a recent study showed that co-infection with an iflavirus increases the insecticidal properties of *Spodoptera exigua* multiple nucleopolyhedrovirus (SeMNPV) occlusion bodies (Carballo et al., 2017; Jakubowska et al., 2016). Insect specific small RNA viruses can also prevent the replication of other viruses in the body of host insects. *In vitro* tests using mosquito-derived C6/36 cells have shown that persistent Palm Creek virus infection (an insect specific flaviviruses) suppresses replication of West Nile Virus and Murray Valley encephalitis virus (Hobson-Peters et al., 2013). This effect is not limited to insects. Harvey et al. (2019) also found that presence of a novel picorna-like virus (Burpengary virus) is negatively associated with Chlamydial disease in the Koala (Harvey et al., 2019).

Several members of these insect small RNA virus families cause clear signs of disease in their hosts, such as infectious flacherie virus in silkworms (Valles et al., 2017), and deformed wing virus (Wilfert et al., 2016) and slow bee paralysis virus in honeybees (Bonning and Miller, 2010). It has been suggested that the picorna-like viruses or other members of Picornavirales can be maintained in insects by horizontal and vertical transmission (Wu et al., 2019). Possibly, the severity of the disease is related to adaptation of virus to the major form of transmission. Horizontally transmitted viruses usually cause more severe and lethal disease compared to vertically transmitted viruses which can be asymptomatic (Ryabov, 2017). The impact of this novel virus (OrPV1) on *O. rhinoceros* larvae and its mode of transmission are unknown, so further investigations are required to understand their natural incidence in different life stages of their host and their geographical distribution.

Currently, our knowledge of *O. rhinoceros* viruses and their interactions with the hosts is very limited. Fortunately, the advent of NGS technology has created a great opportunity for novel virus discovery and the study of host-pathogen interactions. Novel virus discovery will facilitate further studies on interactions between different haplotypes of *O. rhinoceros* and their associated viruses and determine if such interactions might explain the lack of virulence of Oryctes rhinoceros nudivirus (OrNV) to the *O. rhinoceros* haplotype CRB-G. Future studies are essential to determine the potential of *O. rhinoceros* RNA viruses to be used as biological control agents.

## Acknowledgements

This project was supported by the Australian Centre for International Agricultural Research funding (HORT/2016/185), the University of Queensland (UQECR2057321) and Taiwan Ministry of Science and Technology (MOST 106–2311-B-002-002-MY3). The funding source had no role in the study.

## References

Bedford, G.O., 2013. Biology and Management of Palm Dynastid Beetles: Recent Advances. In: Berenbaum, M.R. (Ed.), Annual Review of Entomology, Vol. 58. Annual Reviews, Palo Alto, pp. 353–372.

Bonning, B.C., Miller, W.A., 2010. Dicistroviruses. Annual Review of Entomology 55, 129–150.

Carballo, A., Murillo, R., Jakubowska, A., Herrero, S., Williams, T., Caballero, P., 2017. Co-infection with iflaviruses influences the insecticidal properties of Spodoptera exigua multiple nucleopolyhedrovirus occlusion bodies: Implications for the production and biosecurity of baculovirus insecticides. Plos One 12(5).

Chen, Y.P., Becnel, J.J., Valles, S.M., 2012. RNA Viruses Infecting Pest Insects, 133–170 pp. Insect Pathology, 2nd Edition, edited by

Vega, F.E., Kaya, H.K. Etebari, K., Filipovic, I., Rasic, G., Devine, G.J., Tsatsia, H., Furlong, M.J., 2020. Complete genome sequence of Oryctes rhinoceros nudivirus isolated from the coconut rhinoceros beetle in Solomon Islands. Virus research 278, 197864–197864.

Harvey, E., Madden, D., Polkinghorne, A., Holmes, E.C., 2019. Identification of a novel Picorna-like virus, Burpengary Virus, that is negatively associated with Chlamydial disease in the Koala. Viruses 11(3).

Hobson-Peters, J., Yam, A.W.Y., Lu, J.W.F., Setoh, Y.X., May, F.J., Kurucz, N., Walsh, S., Prow, N.A., Davis, S.S., Weir, R., Melville, L., Hunt, N., Webb, R.I., Blitvich, B.J., Whelan, P., Hall, R.A., 2013. A new Insect-Specific Flavivirus from Northern Australia suppresses replication of West Nile Virus and Murray Valley Encephalitis Virus in co-infected mosquito cells. PloS One 8(2).

Huger, A.M., 2005. The *Oryctes* virus: Its detection, identification, and implementation in biological control of the coconut palm rhinoceros beetle, *Oryctes rhinoceros* (Coleoptera: Scarabaeidae). Journal of Invertebrate Pathology 89(1), 78–84.

Jakubowska, A.K., Murillo, R., Carballo, A., Williams, T., van Lent, J.W.M., Caballero, P., Herrero, S., 2016. Iflavirus increases its infectivity and physical stability in association with baculovirus. Peerj 4.

Le Gall, O., Christian, P., Fauquet, C.M., King, A.M.Q., Knowles, N.J., Nakashima, N., Stanway, G., Gorbalenya, A.E., 2008. Picornavirales, a proposed order of positive-sense single-stranded RNA viruses with a pseudo-T=3 virion architecture. Archives of Virology 153(4), 715–727.

Marshall, S.D.G., Moore, A., Vaqalo, M., Noble, A., Jackson, T.A., 2017. A new haplotype of the coconut rhinoceros beetle, *Oryctes rhinoceros*, has escaped biological control by *Oryctes rhinoceros* nudivirus and is invading Pacific Islands. Journal of Invertebrate Pathology 149, 127–134.

Reil, J.B., Doorenweerd, C., San Jose, M., Sim, S.B., Geib, S.M., Rubinoff, D., 2018. Transpacific coalescent pathways of coconut rhinoceros beetle biotypes: Resistance to biological control catalyses resurgence of an old pest. Molecular Ecology 27(22), 4459–4474.

Ryabov, E.V., 2017. Invertebrate RNA virus diversity from a taxonomic point of view. Journal of Invertebrate Pathology 147, 37–50.

Shelomi, M., Lin, S.S., Liu, L.Y., 2019. Transcriptome and microbiome of coconut rhinoceros beetle (*Oryctes rhinoceros*) larvae. BMC Genomics 20(1).

Valles, S.M., Chen, Y., Firth, A.E., Guerin, D.M.A., Hashimoto, Y., Herrero, S., de Miranda, J.R., Ryabov, E., Consortium, I.R., 2017. ICTV Virus Taxonomy Profile: Iflaviridae. Journal of General Virology 98(4), 527–528.

Valles, S.M., Hashimoto, Y., 2009. Isolation and characterization of Solenopsis invicta virus 3, a new positive-strand RNA virus infecting the red imported fire ant, *Solenopsis invicta*. Virology 388(2), 354–361.

Wilfert, L., Long, G., Leggett, H.C., Schmid-Hempel, P., Butlin, R., Martin, S.J.M., Boots, M., 2016. Deformed wing virus is a recent global epidemic in honeybees driven by Varroa mites. Science 351(6273), 594–597.

Wu, N., Zhang, P.P., Liu, W.W., Cao, M.J., Massart, S., Wang, X.F., 2019. Complete genome sequence and characterization of a new iflavirus from the small brown planthopper (*Laodelphax striatellus*). Virus Research 272.

Yang, X.M., Xu, P.J., Yuan, H., Graham, R.I., Wilson, K., Wu, K.M., 2019. Discovery and characterization of a novel picorna-like RNA virus in the cotton bollworm *Helicoverpa armigera*. Journal of Invertebrate Pathology 160, 1–7.

Zell, R., 2018. Picornaviridae-the ever-growing virus family. Archives of Virology 163(2), 299–317.

